# *Rice yellow mottle virus* is a suitable amplicon vector an efficient production of an anti-leishmianiasis vaccine in *Nicotiana benthamiana* leaves

**DOI:** 10.1101/2023.08.29.555272

**Authors:** PKA Bamogo, F Tiendrébéogo, C Brugidou, D Sérémé, FW Djigma, J Simporé, S Lacombe

**Affiliations:** PHIM Plant Health Institute Montpellier, Univ Montpellier, IRD, CIRAD, INRAE, Institut Agro, Montpellier, France; Institut de L’Environnement et de Recherches Agricoles (INERA), LMI Patho-Bios / Laboratoire de Virologie et de Biotechnologies Végétales, Ouagadougou, Burkina Faso; Université Joseph Ki-Zerbo, Laboratoire de biologie moléculaire et de Génétique (LABIOGENE), Ecole Doctorale Sciences et Technologie, Centre de recherche biomoléculaire Piétro Annigoni (CERBA), Ouagadougou, Burkina Faso

**Keywords:** PSA, RYMV, plant-based, viral vector, leishmaniosis

## Abstract

The suitability of rice yellow mottle virus RYMV as a gene expression vector in plant was assessed using a construct carrying promastigote surface antigen (PSA) C-terminal coding sequence of the parasite protozoan Leishmania. RYMV ORF1 encoding P1 protein has been deleted from the RYMV native genome. The C-terminal PSA gene was substituted for the viral coat protein. PSA is present at the surface of the parasite and displays vaccine properties against canine and human leishmaniosis. RYMV-based vector allowed PSA expression in *Nicotiana benthamiana*. Q-pcr analysis showed that chimeric RYMV carrying PSA gene is able to replicate in *N. benthamiana* leaves. P19 silencing suppressor in combination with the lacked viral vector ORF encoding RYMV Coat Protein (CP) enhanced significantly RYMV tool replication in *N. benthamiana*. RYMV CP played a key role on viral RNA stabilization and acts as a weak silencing suppressor.

The original RYMV-based expression vector allowed PSA protein expression enhancement in N*. benthamiana* without any symptoms. RYMV-based vector could be suitable for functional genomic studies in monocots by VIGS (Viral Induced Gene Silencing) technology.

## Introduction

The use of plants as bioreactor for the production of proteins of interest such as therapeutic or industrial proteins has been booming this last decades. An increasing variety of complex and valuable molecules of interest is produced today in plants bioreactors (see Lee et al 2023 for review). Compared to traditional biological “factories” that are mainly mammalian and microbial, plants bioreactors cannot be infected by mammalian pathogens. Moreover, these alternative systems are cost-effective and easy to scale up. Using *Agrobacterium* mediated transient transformation, agroinfiltration, recombinant proteins can be transiently produced in intact leaves in several days (Kapila et al. 1997). *Nicotiana* plant species such as *N. tabaccum* and *N. benthamiana* are preferentially used due to their high biomass, their fast grow rate and their ability to be stably and transiently transformed via *Agrobacterium* (Conley et al. 2011, Lee et al 2023, Shanmugaral et al. 2020). One of the major constrains in these transient gene expression plant based systems is the relatively limited product yield. Indeed, setting up production systems leading to expression level that is acceptable for economic production (> 50 mg/kg for antibodies) are not trivial process. One of the explanations is the post-transcriptional RNA silencing that the plant cells set up in response to the introduction of foreign nucleic acids (Johansen and Carrington, 2001). The co-expression of viral proteins displaying suppression of RNA silencing activity overcome this limitation and allows increased level of transient expression (Lacombe et al. 2018, Voinnet et al. 2003). Another effective advance in yield improvement has been also based on viruses. It consists on the development of plant viral vectors expressing proteins of interest in plants: amplicon technology. Once introduced in a host plant cells by agroinfiltration, a virus vector engineered to contain a gene of interest replicate and the protein of interest can be produced rapidly in significant quantities (Gelba et al. 2007; Lico and al. 2008). This method has been shown to work with numerous proteins. Viral vectors used so far have been mostly derived from *Tobacco mosaic virus* (TMV) and *Potato virus X* (PVX) combined with host plants such as *N.tabaccum* and *N. benthamiana* (Daniell et al. 2009; Marillonnet et al. 2005; Komarova et al. 2006). First generation of viral vectors were based on the addition of the gene of interest into complete viral genome. However, a negative correlation was demonstrated between insert size and virus stability due to the rapid loss of the transgene during viral replication (Avesani et al. 2007; Gelba et al. 2004). A more efficient generation of viral vectors were based on deconstruct virus genome where only elements required for replication were maintained. This allows the insertion of large gene of interest while maintaining viral genome size and stability (Gelba et al. 2004, 2005).

*Rice yellow mottle virus* is a member of sobemoviruses genus. It genome is a single-strand positive-sense RNA molecule of around 4.5 kb. Its 5’ end has a covalently linked viral protein (VPg) and its 3’ end is not polyadenylated (Tamm and Truve 2000; Ngon et al. 1994; Yassy et al. 1998). The genome contains five protein coding ORFs (Fig 1). ORF1, x, 2a and 2b are translated from the genomic RNA whereas ORF3 is translated from the sub-genomic RNA (Ling et al. 2013). Except for ORFx that has been lately identified (Ling et al. 2013), functions of all the other ORFs have been well characterized. ORF1 encodes P1 protein that has been described as a RNA silencing suppressor (Voinnet et al. 1999, Siré et al. 2008, Lacombe et al. 2010). P1 is also involved in virus movement (Bonneau et al. 1998). ORF2a and 2b encode polyproteins that are cleaved to produce the VPg, a serine protease, a helicase and a RNA-dependent RNA polymerase (RdRp) (Tamm and Truve 2000; Ngon et al. 1994). ORF3 encodes for the Coat Protein (CP) (Tamm and Truve 2000). Functional studies demonstrated that both P1 and CP are dispensable for virus replication in rice protoplast (Bonneau et al. 1998, Brugidou et al.1995). Using the RYMV infectious cDNA clone developed by Brugidou et al. (1995), P1 non-requirement for viral replication was confirmed into rice plants (Nummert et al. 2017). RYMV natural hosts are restricted several species of the genera *Oryza* (Fauquet and Thouvenel, 1987). It is mechanically transmissible to rice but also to several graminaceous species. RYMV dilution end-point quite low (10^-11^) reveals important virus titer in infected samples suggesting strong virus stability and high replication rate (Fauquet and Thouvenel, 1987). Using the RYMV infectious cDNA clone, viral replication was not only detect in rice plants but also in wheat, oat and barley monocotyledons plants and in *Arabidopsis thaliana* and *N. benthamiana* dicotyledons plants (Nummert et al. 2017). Based on its relatively simple genome well characterized, the unnecessary P1 and CP for viral replication, its high replication rate and its ability to replicate in *N. benthamiana*, RYMV represent a candidate of choice for the development of a new amplicon tool for the production of proteins of interest in *N. benthamiana*.

**Figure 1:**
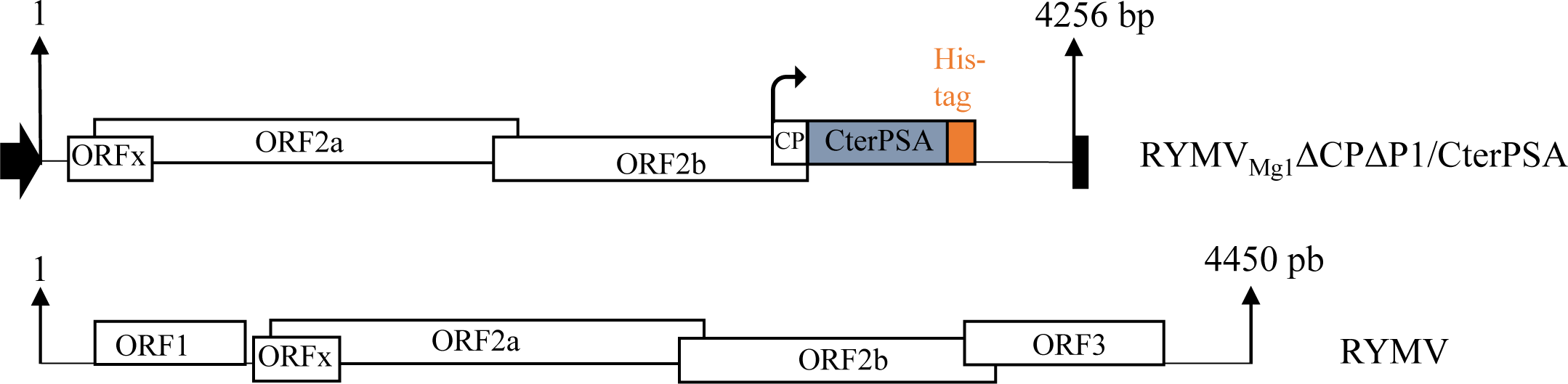
Schematic representation of RYMV_Mg1_ΔCPΔP1/CterPSA construct based on RYMV genome. For both RYMV_Mg1_ΔCPΔP1/CterPSA construct and RYMV reference, ORFs are represented by white boxes. Sizes of both sequence are indicated. For RYMV_Mg1_ΔCPΔP1/CterPSA construct, the insert encoding for the C-terminal part of *Leishmania infantum* PSA (CterPSA) is indicated by the blue box. An His tag represented by the orange box is fused to its C-terminal part. Black arrow and black box represent CaMV duplicated 35S promoter and CaMV polyA signal sequence / terminator, respectively.

Here, we addressed this point by developing an RYMV amplicon vector in which P1 and CP genomic sequences non-necessary for virus replication were removed. We focused on Promastigote Surface Antigen (PSA) protein as proteins of interest to produce. Indeed, this protein coming from the parasitic protozoan *Leishmania infantum* has been characterized as displaying efficient vaccine properties against canine leishmaniasis (Lemesre et al. 2005, 2007). Moreover, we previously demonstrated that an active PSA protein can transiently accumulated in *N. benthamiana* leaves after agroinfiltration of a PSA coding sequence driven by the 35S constitutive promoter. However, despite optimized RNA silencing suppression and subcellular localization, PSA accumulation only reached 0.4 mg/kg of fresh weigh which is insufficient for acceptable for economic production (Lacombe et al. 2018). Here, we replaced the CP subgenomic coding sequence by the PSA sequence into RYMV amplicon vector. Series of experiments were performed to optimize RYMV amplicon vector replication in *N. benthamiana* leaves allowing an improved protein of interest accumulation. This work demonstrates that RYMV amplicon tool and procedure developed here are effective for transient accumulation of proteins of interest in *N. benthamiana* in a rapid, efficient and low-cost way.

## Results

### Design of RYMV amplicon vector

Construction of amplicon tools has to minimize genome structure modification in order to avoid the expulsion of sequences encoding protein of interest during viral replication. Consequently, sequences of interest have to be inserted instead of viral sequences non-necessary for viral replication. Previous works demonstrated that both P1 protein encoded by ORF1 and CP encoded by ORF3 were dispensable for virus replication in rice protoplasts and plants (Bonneau et al. 1998, Brugidou et al. 1995, Nummert et al. 2017). Both P1 and CP sequences were removed and sequence of interest has been inserted instead of the CP ORF into the highly replicative subgenomic part of the RYMV genome (Figure 1). CP ORF 5’ extremity overlaps with ORF2b. To maintain ORF2b coding sequence integrity, CP ORF replacement has to be partial maintaining the ORF2b / CP ORF overlapping sequence. Consequently, a fusion protein CP-protein of interest is expected to be produced. P1 protein has been previously described as a suppressor of the RNA silencing defense mechanism (Lacombe et al. 2010, Siré et al. 2008). It has been shown that the efficiency of RNA silencing suppression varies according to RYMV isolates from weak to strong suppressors such as P1 from Madagascar isolate (Mg1) and P1 from Tanzania isolate (Tz3), respectively (Lacombe et al. 2010, Siré et al. 2008). Consequently, we chose to consider Mg1 genetic background for the amplicon tool developed here as only a weak RNA silencing suppression activity would be missing (Figure 1). As protein of interest, the promastigote surface antigen (PSA) protein from the parasitic protozoan *Leishmania infantum* displaying anti-leishmaniasis vaccine properties has been considerate. We previously managed to produce this protein in *N. benthamiana* using 35S expression vector (35S::PSA) but failed to reach an acceptable production yield (Lacombe et al. 2018). Here, we considered only the carboxy terminal part of the PSA (CterPSA) as it induced the higher level of protection in tested dogs (Petitdidier et al. 2016) and produced the resulting amplicon vector RYMV_Mg1_ΔCPΔP1/CterPSA (Figure 1).

### RYMV-based amplicon RNA accumulated efficiently in *Nicotiana benthamiana* with CP *trans* complementation

Under natural conditions, RYMV hosts are restricted to grasses species such rice. We first addressed whether RYMV_Mg1_ΔCPΔP1/CterPSA RNA accumulated in *N. benthamiana* leaves. We used an optimized RNA silencing suppression condition that has been previously demonstrate to favor mRNA accumulation of gene of interest in transient *N. benthamiana* assays (Lacombe et al. 2018). We co-infiltrated *N. benthamiana* with a 1:1 ratio of RYMV_Mg1_ΔCPΔP1/CterPSA construct and the cocktail of RNA silencing suppressors described in Lacombe et al. 2018. The cocktail optimizes the RNA silencing suppression. It consists of 1:1:1 combination of three viral suppressors (P0 from *Beet western yellows virus*, P1 from RYMV and P19 from *Cymbidium ring spot virus*) acting on different steps of RNA silencing pathway. The empty vector was used as negative controls. Infiltrated leaves were harvested four days post infiltration (4 dpi) for total RNA extraction followed by semi quantitative RT PCR of the CterPSA sequence. As expected, no amplification was observed for the empty vector control samples. A low amplification was observed for the RYMV_Mg1_ΔCPΔP1/CterPSA samples (Figure 2). This indicated that RYMVΔCPΔP1/CterPSA construct can be transcribed when expressed in *N. benthamiana* leave but the corresponding RNA accumulated poorly even under optimized RNA silencing suppression condition.

**Figure 2:**
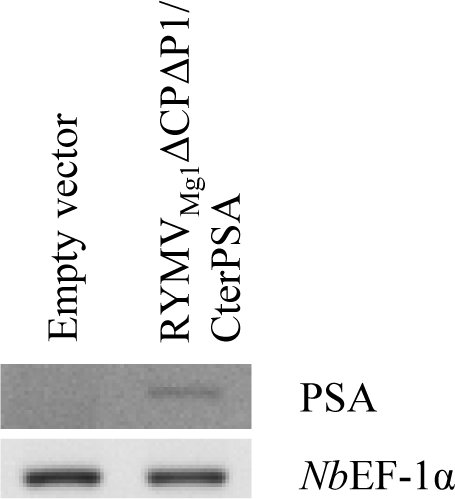
RYMV_Mg1_ΔCPΔP1/CterPSA RNA accumulation in *Nicotiana benthamiana* leaves. Semi-quantitative RT-PCR were performed on RNA extracted from *N. benthamiana* leaves infiltrated with either empty vector or RYMV_Mg1_ΔCPΔP1/CterPSA construct together with the cocktail of RNA silencing suppressors. Reverse transcriptions were performed using a mix of oligodT and CterPSA specific primers. PCR amplifications were performed using CterPSA specific primers. *Nb*EF1-α expression was used as internal control. Three independent experiments were performed. Results presented here are representative of the three experiments.

P1 and CP have been described as dispensable for viral replication in rice protoplasts and plants. However, we cannot exclude that they could be required for an efficient replication of RYMV_Mg1_ΔCPΔP1/CterPSA amplicon in *N. benthamiana* leaves. It does not seem to be the case for P1 as its presence in the RNA silencing suppressor cocktail used previously led only to a weak RYMV_Mg1_ΔCPΔP1/CterPSA RNA accumulation. To test the effect of CP *trans* complementation, 35S-driven CP construct was co-infiltrated together with RYMV_Mg1_ΔCPΔP1/CterPSA in *N. benthamiana* leaves with RYMV_Mg1_ΔCPΔP1/CterPSA:CP:empty vector 1:0,5:0,5 ratio. CP RNA and protein accumulations were evaluated by semi quantitative RT-PCR and western blot, respectively (Figure 3 A, 3B). Signals were detected at both RNA and protein levels but quite weakly. As CP is expressed as exogenous gene, its expression should be affect by the RNA silencing defense pathway. To test this, RYMVΔCPΔP1/CterPSA and CP constructs were co-inoculated with P19 RNA silencing suppressor in a 1:0.5:0.5 ratio (amplicon:CP:P19). Semi quantitative RT-PCRs and western blots show that the presence of P19 improves both CP RNA and protein accumulations indicating that P19 protected CP from RNA silencing degradation (Figure 3A, 3B). RYMV_Mg1_ΔCPΔP1/CterPSA RNA accumulations were followed by qRT-PCR for all tested conditions with or without CP and/or P19 (Figure 3C). Improved amplicon RNA accumulation was only observed when both CP and P19 were present. This suggests that improved CP accumulation due to P19 increases by *trans* complementation RYMV_Mg1_ΔCPΔP1/CterPSA RNA accumulation in *N. benthamiana*.

**Figure 3:**
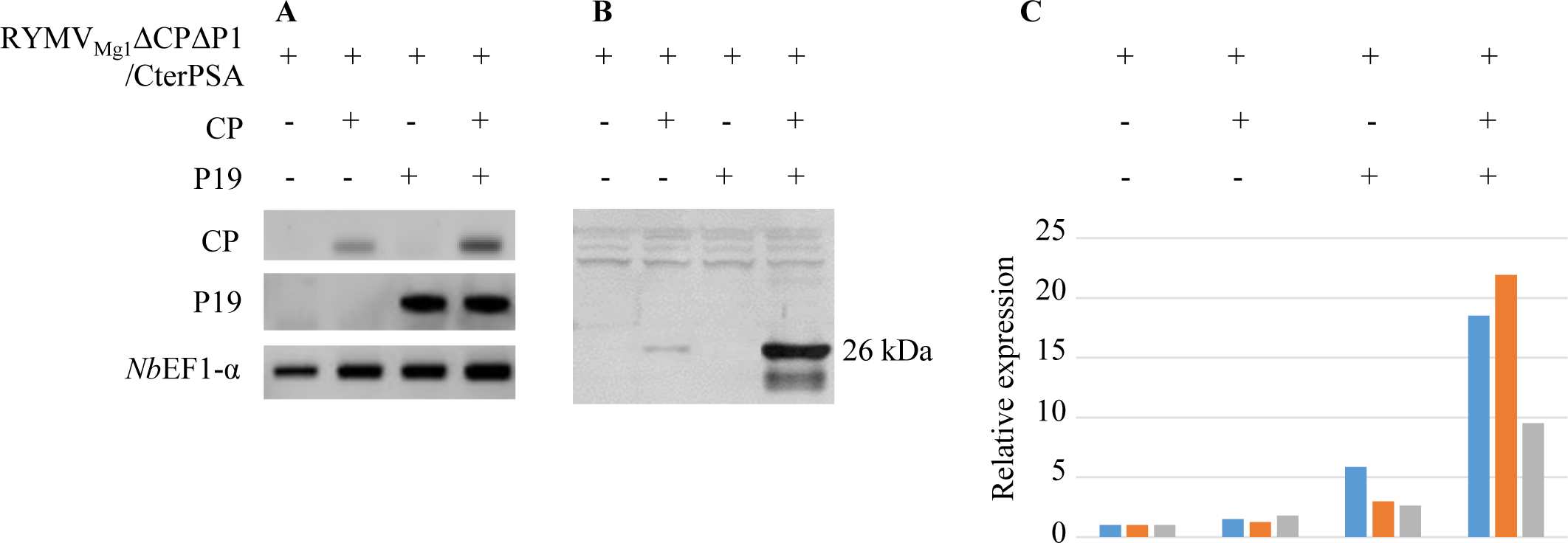
P19 RNA silencing suppressor and CP act synergistically to enhance RYMV_Mg1_ΔCPΔP1/CterPSA RNA accumulation. *N. benthamiana* leaves were infiltrated with RYMV_Mg1_ΔCPΔP1/CterPSA construct on its one (RYMV_Mg1_ΔCPΔP1/CterPSA : empty vector 1:1 ratio) or with CP and P19 constructs either alone (RYMV_Mg1_ΔCPΔP1/CterPSA:CP or P19:empty vector 1:0,5:0,5 ratio) or together (RYMV_Mg1_ΔCPΔP1/CterPSA:CP:P19 1:0,5:0,5 ratio). Infiltrated leaves were harvested at 5 dpi. Three independent experiments were performed. **A-** Semi-quantitative RT-PCR were performed after RNA reverse transcription performed with a mix of oligodT and CterPSA specific primers. PCR amplifications were performed using CP and P19 specific primers. *Nb*EF1-α expression was used as internal control. Results presented here are representative of the three experiments. **B-** Total soluble proteins were prepared from same *N. benthamiana* samples. Equal among of proteins (10 µg) was separed by SDS-PAGE and analysed by western immunoblotting using anti-CP antibody. Size in kDa of the CP signals is indicated. Results presented here are representative of the three experiments. **C-** RYMV_Mg1_ΔCPΔP1/CterPSA RNA accumulation was evaluated by q-RTPCR for cDNA samples prepared in A. CterPSA specific primers were used. Normalizations were done using GAPDH used as housekeeping internal control. Expressions were evaluated relatively to the condition “RYMV_Mg1_ΔCPΔP1/CterPSA alone”. The three independent experiments are presented by distinct colors.

### Optimization of RYMV_Mg1_ΔCPΔP1/CterPSA RNA accumulation in *N. benthamiana*

To evaluate if an increased quantity of CP and P19 could improve their positive effect on RYMV_Mg1_ΔCPΔP1/CterPSA RNA accumulation, the initial 1:0.5:0.5 (amplicon:CP:P19) ratio was compared with three other ratios with increased quantities of CP and P19 constructs (Figure 4). RYMV_Mg1_ΔCPΔP1/CterPSA RNA accumulations were followed by qRT-PCR. Results demonstrated that increasing CP and P19 quantity up to 1:2.5:5 ratio (amplicon:CP:P19) improved RYMV_Mg1_ΔCPΔP1/CterPSA RNA accumulation compared to the initial 1:0.5:0.5 condition. Beyond this threshold, a 1:4:5 ratio has a negative effect with a reduction of RYMV_Mg1_ΔCPΔP1/CterPSA RNA accumulation compared to the initial 1:0.5:0.5 condition (Figure 4). Consequently, we retained 1:2.5:5 ratio (amplicon:CP:P19) has the optimal ratio for amplicon RNA accumulation in *N. benthamiana* leaves.

**Figure 4:**
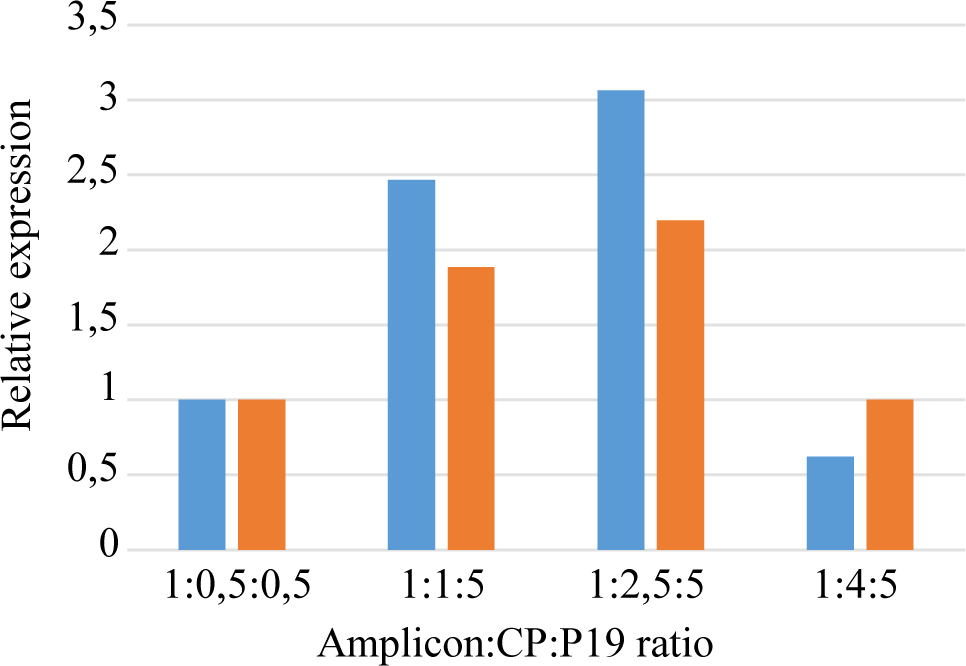
Optimal CP:P19 ratio for RYMV_Mg1_ΔCPΔP1/CterPSA RNA accumulation. *N. benthamiana* leaves were infiltrated with RYMV_Mg1_ΔCPΔP1/CterPSA construct together with CP and P19 constructs. Four different amplicon:P19:CP ratio were considerate. Infiltrated leaves were harvested at 5 dpi. Two independent experiments were performed. q-RT-PCR were performed after RNA reverse transcription using a mix of oligodT and CterPSA specific primers. PCR amplifications were performed using CterPSA specific primers. GAPDH was used as housekeeping internal control. For each ratio, expressions were evaluated relatively to the initial 1:0,5:0,5 ratio. The two independent experiments are presented by distinct colors.

Once present in host cells, viruses multiply and move from cell to cell to invade systemic tissues. In order to determine tissues with the best amplicon RNA accumulation, we evaluated the ability of RYMV_Mg1_ΔCPΔP1/CterPSA amplicon to move through *N. benthaniama* plants under the optimal 1:2.5:5 ratio (amplicon:CP:P19) conditions. Infiltrated (1), adjacent (2) and systemic (3) tissues were considerate to evaluate amplicon RNA accumulation by semi quantitative RT-PCR (Figure 5A). For both 35S::PSA control and RYMV_Mg1_ΔCPΔP1/CterPSA constructs, PSA amplifications were detected only in infiltrated areas indicating that RYMV_Mg1_ΔCPΔP1/CterPSA construct is unable to move from cell to cell. Then, we followed by q-RT-PCR PSA RNA accumulation over the time to determine stages displaying the best RNA accumulation (Figure 5B, 5C). Both RYMV_Mg1_ΔCPΔP1/CterPSA repeats showed a faint accumulation until 9 dpi and them a strong increase until 12 dpi followed by a decrease (Figure 5B). The 35S::PSA control construct acted differently with an important accumulation induction since 2 dpi to 5 – 7 dpi and them a stabilization (Figure 5C). Based on these results, we retained infected areas at 12 dpi with 1:2.5:5 ratio (amplicon:CP:P19) as conditions displaying optimal RYMV_Mg1_ΔCPΔP1/CterPSA RNA accumulation.

**Figure 5:**
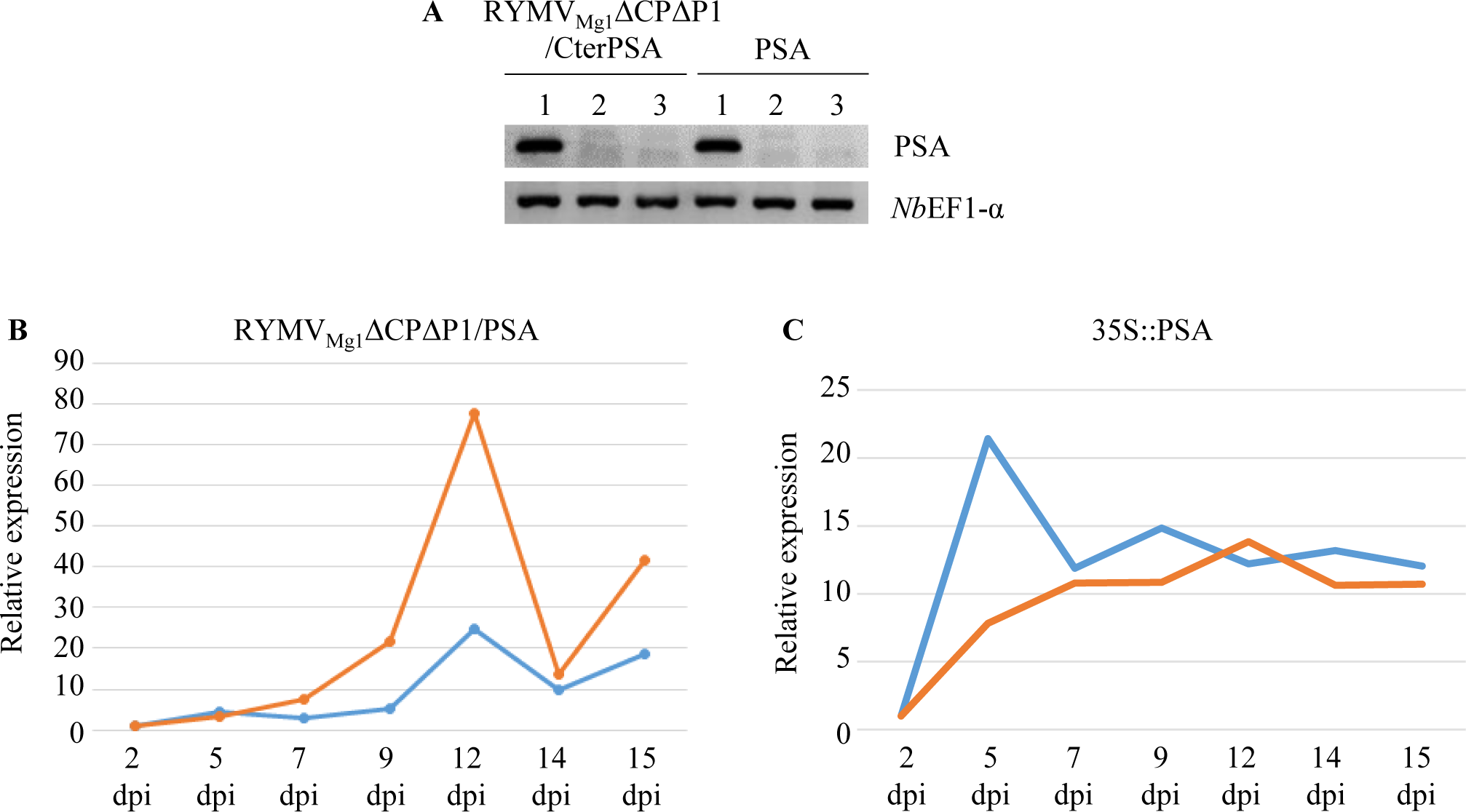
RYMV_Mg1_ΔCPΔP1/CterPSA RNA spacio-temporal accumulation pattern. *N. benthamiana* leaves were infiltrated with RYMV_Mg1_ΔCPΔP1/CterPSA construct together with CP and P19 constructs under the optimal 1:2,5:5 amplicon:CP:P19 ratio. 35S::PSA construct in combination with P19/P1/P0 RNA silencing suppressor cocktail was used as control. Two independent experiments were performed. **A-** At 5 dpi, RNAs were prepared from infiltrated area (1), adjacent non infiltrated area of the same leaves (2) and systemic leaves (3). Semi-quantitative RT-PCR were performed after reverse transcriptions using a mix of oligodT and CterPSA specific primers. PCR amplifications were performed using CterPSA specific primers. *Nb*EF1-α expression was used as internal control. For both RYMV_Mg1_ΔCPΔP1/CterPSA RNA (**B**) 35S::PSA (**C**) constructs, RNA accumulation was followed from 2 to 15 days-post-inoculation (dpi) in infiltrated area. q-RT-PCR were performed after RNA reverse transcription using a mix of oligodT and CterPSA specific primers. PCR amplifications were performed using CterPSA specific primers. Normalizations were done using GAPDH used as housekeeping internal control. . Expressions were evaluated relatively to the 2 dpi time point. The two independent experiments are presented by distinct colors.

### RYMV_Mg1_ΔCPΔP1/CterPSA autonomously replicated in *N. benthamiana* leaves

Once introduced into plant cells, amplicon construct under constitutive promoter is expected to be first host cell nuclear transcribed as mRNA to produce viral proteins required for viral replication such as the viral RdRP. Them, autonomous viral replication can occur to amplify viral RNA through the synthesis of double strand RNA mediated by viral RdRP. Consequently, to demonstrate that RYMV_Mg1_ΔCPΔP1/CterPSA autonomous replication occurred, we searched for antisens RNA accumulation in *N. benthamiana* leaves infiltrated by RYMV_Mg1_ΔCPΔP1/CterPSA construct under the optimal conditions previously determined. 35S::PSA construct was used as negative control. To detect antisens amplicon RNA, reverse transcriptions were performed using PSA primer hybridizing the negative strand RNA (PSA_for_) followed by PCR amplification with PSA specific primers (Figure 6). PCR amplifications were detected for all RYMV_Mg1_ΔCPΔP1/CterPSA samples. Surprisingly, amplifications were also detected in 35S::PSA negative control samples suggesting a possible DNA contamination. However, no amplifications occurred using *Nb*EF1-α primers on these antisens cDNA samples whereas these *Nb*EF1-α primers allowed amplifications on cDNA populations produced using oligodT primer (Figure 6). Consequently, we excluded that PSA amplification could be due to DNA contaminations. Once introduced in *N. benthamiana* cells, both RYMV_Mg1_ΔCPΔP1/CterPSA and 35S::PSA constructs are recognized as foreign nucleic acid and induced RNA silencing defense mechanism. During this process, antisens RNA are produced from foreign RNA by an endogenous RdRP. Double strand RNA are then processed by Dicer enzyme to produce 21, 24 nt siRNA (Johansen and Carrington, 2001). RNA silencing suppressor used in our experiments (P19, P1 and P0) acts downstream the double strand RNA step. That could explain PSA PCR amplification in antisens cDNA samples for 35S::PSA negative control. Consequently, in our transient expression system, detection of antisens RNA is not a proof of RYMV_Mg1_ΔCPΔP1/CterPSA autonomous replication as it can be produced by either viral or endogenous RpRP.

**Figure 6:**
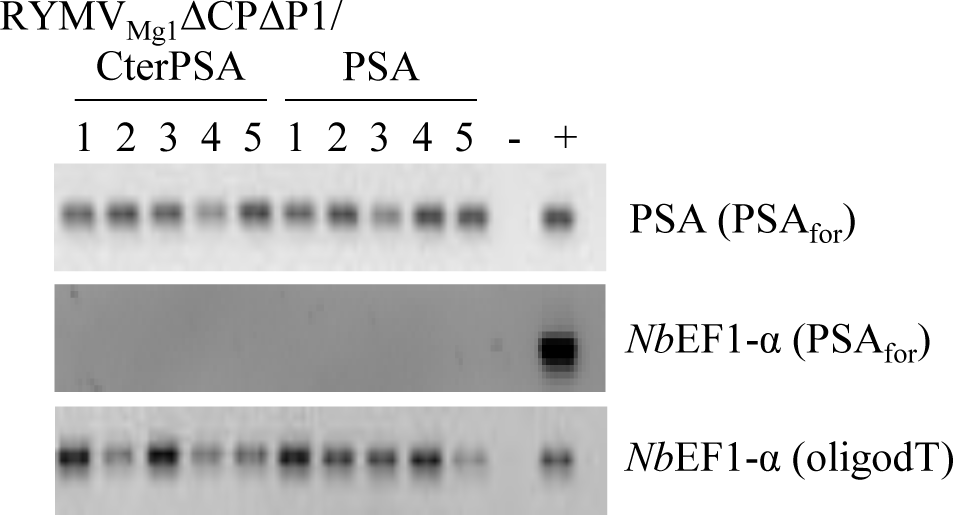
RYMV_Mg1_ΔCPΔP1/CterPSA minus strand RNA accumulation. *N. benthamiana* leaves were infiltrated with RYMV_Mg1_ΔCPΔP1/CterPSA construct together with CP and P19 constructs under the optimal 1:5:2,5 amplicon:P19:CP ratio. 35S::PSA construct in combination with P19/P1/P0 RNA silencing suppressor cocktail was used as control. Five individual plants, noted from 1 to 5 were considerate from both constructs. RNA were prepared from infiltrated leaves at 7 dpi. Reverse transcription were done using either PSA primer hybridizing minus strand of Cterm part of the PSA sequence (PSA_for_) or oligodT primer. Primer used for reverse transcription is noted in brackets. PCR amplifications were performed using PSA specific primers or *Nb*EF1-α primers. Negative (-) and positive (+) PCR controls were used.

It has been shown that genome of viruses such as RYMV belonging to sobemovirus is a single strand RNA with non-polyadenylated (non-polyA) 3’ end (Yassy et al. 1988). Consequently, if RYMV_Mg1_ΔCPΔP1/CterPSA amplicon is active, a mix of polyA and non-polyA RNA coming from host cell nuclear transcription driven by the constitutive promoter and from autonomous viral replication respectively, should accumulate in infiltrated *N. benthamiana* cells. To evaluate if non-polyA RNA accumulated, two cDNA populations were done. The first one produced by reverse transcriptions with oligodT primer represented polyA mRNA population only. The second one produced by reverse transcriptions with a mix of oligodT and CterPSA specific primers (CterPSA_spe_) represented both polyA mRNA and non-polyA RNA populations if present. q-RT-PCR on CterPSA region were performed on these two cDNA populations for both RYMV_Mg1_ΔCPΔP1/CterPSA and 35S::PSA samples previously obtained. Expressions for cDNA populations representing polyA plus non-polyA RNA were evaluated relatively to the expression for cDNA population representing only polyA mRNA (Figure 7A, 7B). For both RYMV_Mg1_ΔCPΔP1/CterPSA and 35S::PSA samples, q-RT-PCR signals were stronger in polyA plus non-polyA cDNA populations compared to polyA ones. In the case of 35S::PSA samples, these increased q-RT-PCR signals could not be explained by an accumulation of non-polyA RNA but rather by an improved reverse transcription efficiency with CterPSA_spe_ primer compared to oligodT. In the case of RYMV_Mg1_ΔCPΔP1/CterPSA samples, this increased amplification is significantly stronger than in the case of 35S::PSA samples (Figure 7C). This suggests that it cannot be only due to the improved RT efficiency with CterPSA_spe_ primer but also to an accumulation of non-polyA RNA characterizing an effective autonomous RYMV_Mg1_ΔCPΔP1/CterPSA replication.

**Figure 7:**
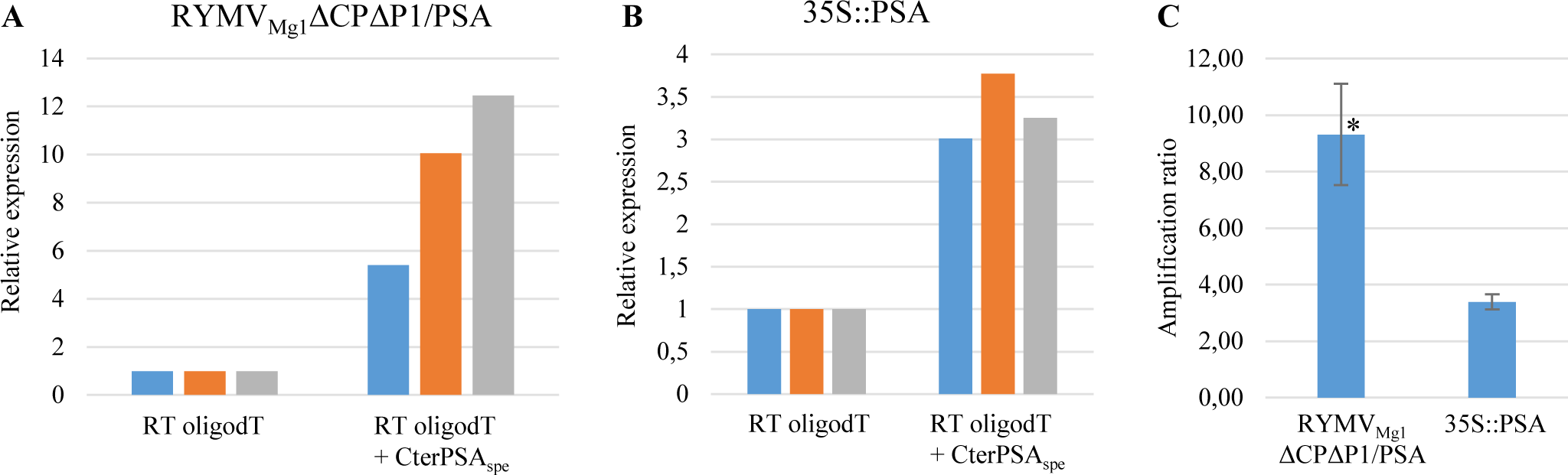
Comparison of total RNA accumulation versus polyA mRNA accumulation. Total RNA prepared previously from *N. benthamiana* leaves infiltrated either with RYMV_Mg1_ΔCPΔP1/CterPSA or 35S::PSA constructs was reverse transcribed using either oligodT or a mix of oligodT and CterPSA specific primer (CterPSA_spe_). Three out of the five individual plants were considerate. For both RYMV_Mg1_ΔCPΔP1/CterPSA **(A)** and 35S::PSA **(B)** samples, q-RT-PCR were performed using CterPSA specific primers. Normalizations were done using GAPDH used as housekeeping internal control. For each construct, expressions were evaluated relatively to the initial RT oligodT condition. The three independent plants are presented by distinct colors. **C-** Ratio of total RNA versus mRNA accumulations were compared between RYMV_Mg1_ΔCPΔP1/CterPSA and 35S::PSA samples. * Student t-test p=0,001

### RYMV_Mg1_ΔCPΔP1/CterPSA allows an improved CterPSA protein accumulation in infiltrated *N benthamiana* leaves

To evaluate that RYMV_Mg1_ΔCPΔP1/CterPSA construct is able to accumulate the expected protein of interest, CterPSA accumulation was evaluated by western blot on *N. benthamiana* samples infiltrated with RYMV_Mg1_ΔCPΔP1/CterPSA construct. Amplicon construct was infiltrated on its one, with either CP or P19 and with both CP and P19 under the optimal conditions previously established. 35S::PSA construct was used as control as established in Lacombe et al. 2018 (Figure 8). As expected, control samples accumulated 48 kDa PSA protein. For RYMV_Mg1_ΔCPΔP1/CterPSA samples, CterPSA accumulation was only detected in samples expressing both CP and P19 at the expected 21 kDa size. This confirms at protein level the requirement of P19 mediated CP *trans* complementation. Interestingly, the intensity of western signal was stronger for RYMV_Mg1_ΔCPΔP1/CterPSA samples under optimal conditions compared to the 35S::PSA control sample (Figure 8). This reinforces previous finding that demonstrated that RYMV_Mg1_ΔCPΔP1/CterPSA is able to replicate autonomously once introduced into *N.benthamiana* cells under optimal conditions and demonstrated that this active replication greatly improves protein of interest accumulation.

**Figure 8:**
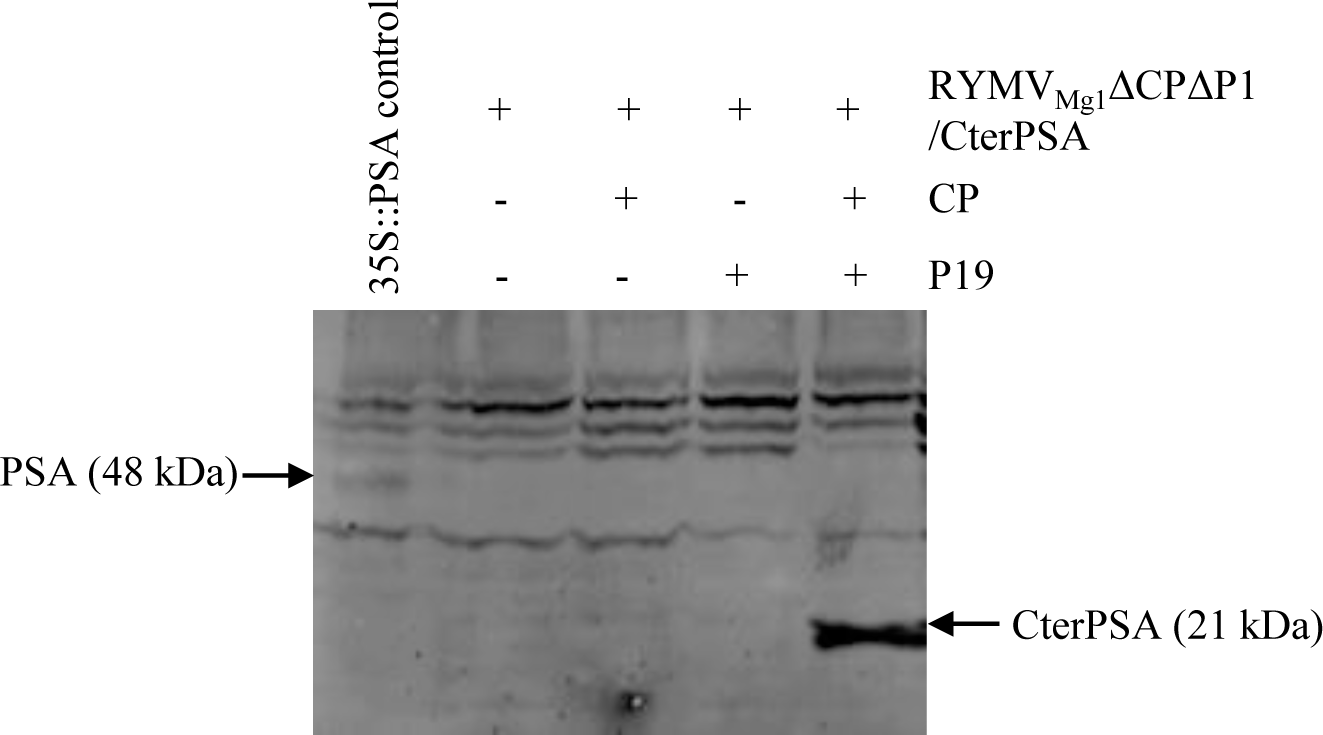
RYMV_Mg1_ΔCPΔP1/CterPSA produces CterPSA protein under optimized condition. *N. benthamiana* leaves were infiltrated with RYMV_Mg1_ΔCPΔP1/CterPSA construct on its one (RYMV_Mg1_ΔCPΔP1/CterPSA : empty vector 1:1 ratio) or with CP and P19 constructs either alone (RYMV_Mg1_ΔCPΔP1/CterPSA:CP or P19:empty vector 1:0,5:0,5 ratio) or together (RYMV_Mg1_ΔCPΔP1/CterPSA:CP:P19 1:2,5:5 ratio). Infiltrated areas were harvested at 12 dpi. 35S::PSA construct in combination with P19/P1/P0 RNA silencing suppressor cocktail was used as control in the same condition as described in Lacombe et al. 2018. Total soluble proteins were prepared. Equal amounts of proteins (100 µg) were separated by SDS-PAGE and analyzed by immunoblotting using an anti-PSA antibody. Bands corresponding to PSA constructs and their corresponding size are indicated. Three independent experiments were performed. Results presented here are representative of the three experiments.

## Discussion

In 1995 a full-length RYMV vector which is able to reproduce the infection when inoculated onto rice plants have been reported on (Brugidou *et al*., 1995). In this study we describe the construction of a 35S-driven new viral expression vector that could efficiently be deliver to *N benthamiana* cells by agroinfection based on RYMV. The vector developed here is a CP deletion mutant of the full-length RYMV vector that efficiently expressed the transgene when complemented with the missing CP in a condition of RNA silencing suppression with P19 this for an accurate ratio of RYMV_Mg1_ΔCPΔP1/CterPSA: CP:P19 of 1:2.5:5. The complementation with only CP did not lead to the accumulation of PSA protein (Figure 8) and RT PCR analysis showed that the CP is strongly silenced when the agroinfection was done without P19 (Figure 3A, 3B). This indicated that CP and P19 interact to significantly enhance RVMV-based vector functioning. Therefore, it is hypothesized that P19 had an indirect effect on RYMV_Mg1_ΔCPΔP1/CterPSA replication it protected the CP that was only able to support RYMV_Mg1_ΔCPΔP1/CterPSA functioning once it is protect by P19 from RNA silencing established by *N benthamiana*. Because the CP was strongly silenced, it is also hypothesized that the RNA sequence of the RYMV CP ORF may be a potent inducer of RNA silencing.

P1 protein is known as a movement protein for RYMV (Bonneau *et al*., 1998; Meier *et al*., 2006; Nummert *et al*., 2017; Sivakumaran *et al*., 1998). Ours vector is also a P1 deletion mutant vector the experiment we performed to assessed ours P1 deletion mutant movement led to the conclusion that RYMV_Mg1_ΔCPΔP1/CterPSA was not able to move either locally or systemically. That results are on the contrast of Nummert et al recent findings about a P1 independent movement of RYMV in host and non-host plant species including *N benthamiana* (Nummert *et al*., 2017). Nummert et al assessed the P1 deletion mutant RYMV movement without a negative control. An efficient experimental procedure should have include a control free from any viral genome sequence for which no movement should be detected locally. In our work RYMV_Mg1_ΔCPΔP1/CterPSA vector behaved the same way as the control PSA, we have not detected any movement outside the infiltration area (Figure 5A). Another point of contradiction between ours findings and Nummert et al work is the detection of the RYMV minus strand as an evidence of replication. Here the negative control which is the PSA sequence free from any viral genome sequence also led to the detection of an apparent minus strand (Figure 6). We hypothesized that this apparent minus strand is originated from the plant RNA silencing pathway when the RdRP build a complementary strand of the foreign RNA for the RNA duplex formation. Thus the detection of minus strand would not be an efficient evidence of viral vector replication. We suggest a statistical student t test on two groups of RNA (mRNA and Total RNA) to evidence the augmentation of total RNA in transformed *N. benthamiana* leaves compare to mRNA in the same transformed leaves due to viral vector replication (Figure 7C).

Since the classical prokaryote *Escherichia coli* bio system failed in accumulating PSA, a standard plant system based on *N. benthamiana* has been optimized for the production of putative active PSA by abolishing the RNA silencing defense mechanism set up by the plant against PSA ectopic expression using three viral suppressors acting at different steps of the RNA silencing pathway combined with the PSA ER retention (Lacombe et al. 2018). PSA protein detected in this study compare to the optimized expression performed according to Lacombe et al was greater (Figure 8). The RYMV-based vector also presented the advantage of a long lasting overexpressing of a protein of medical interest in a short time as PSA protein expression trough this biotechnological tool started 5 five days after infiltration and lasted over 12^th^ day after infiltration (Figure 5). Ours experiments accurately sort out the optimal RYMV_Mg1_ΔCPΔP1/CterPSA:CP:P19 combination and the optimal day of production which were respectively 1:2.5:5 and 12 days post infiltration. However, it is recommended to perform a mass spectrometry analysis to well characterize the PSA Cterm protein produce in our study.

Altogether, these data suggest that the new RYMV viral-based expression vector in *N. benthamiana* transient expression system may be suitable for the rapid, easy and efficient production of valuable proteins of interest.

To prevent or treat infectious diseases especially in situations requiring rapid responses such as pandemic threats and bioterrorism events it is crucial to develop therapeutics such as vaccine (antigenic pathogen peptides) but also prophylactics like monoclonal antibodies (mAbs). As we saw upper, original viral-based biotechnological tools like RYMV based viral vector are useful for antigenic peptides production for vaccine purpose. It could also be useful for neutralizing antibodies production in plant-based biosystem. Antibodies are complex glycoproteins consisting of four polypeptides, linked by disulfide bridges and non-covalent bonds. Despite their complexity, mAbs can be easily expressed in plants biosystems (Fischer *et al*., 2003; Hiatt *et al*., 1989; Ma *et al*., 2005). Rapid transient expression of neutralizing antibody can be easily achieve by expressing a full length antibody through plants co-infecting with two viral vectors harboring the light chain (LC) and the heavy chain (HC) respectively (Alamillo *et al*., 2006; Verch *et al*., 1998). However, the transient expression of heterooligomeric proteins such as mAbs thanks to viral vectors technology presents some drawback when the production is achieved through the co-delivery of viral vectors built on the same virus backbone. In fact, this feature results in early segregation and subsequent preferential amplification of one of the vectors in one cell. This problem is overcome by utilizing two sets of vectors derived from non-competing viruses to express the two different mAb chains (Giritch *et al*., 2006). The two chimeric viruses interact with different host proteins to sustain their replication and movement without interfering with each other. As a result, neither of the two vectors gains a replicative advantage over the other, allowing efficient co-expression of HC and LC in the same cells. The RYMV_Mg1_ΔCPΔP1/CterPSA viral vector developed here could be an additional viral vector for an exploitation in plant biotechnological domain for mAbs production.

## Conclusion

In this study, we reported successful development of a new expression viral vector based on RYMV and using *N. benthamiana* transient expression system. We showed that this RYMV-based vector when complementing with a combination of RNA silencing suppressors P19 and RYMV CP is highly suitable for producing PSA protein. The viral vector was not able to perform movement in transformed *N. benthamiana* plants, it was stable and allowed a long lasting production. This work will serve as bases for further opening out to other proteins of interest such as antibodies.

## Material and methods

### RYMV_Mg1_ΔCPΔP1/ Cter PSA design, synthesis and cloning

#### Expression vectors

The experiments were performed using *Agrobacterium tumefaciens* strains harbouring pCambia A4: RYMV_Mg1_ΔCPΔP1/ Cter PSA. This construct represent the backbone of RYMV virus genome deleted from CP and P1 sequences and harbour the Cterm part of *Leishmania infantum* surface antigen gene sequence which display interesting vaccine properties toward canine leishmaniosis (Petitdidier *et al*., 2016). In addition to the viral expression vector construct (pCambia A4: RYMVMg1ΔCPΔP1 / Cter PSA), we also used a construct expressing the entire protein of PSA gene pCambia A4:PSA (Lacombe *et al*., 2018), a construct expressing the capsid protein (CP) of RYMV (pCambia A4:CP_Mg1_) and three other constructs expressing plant RNA silencing suppressors. These plant RNA silencing suppressors constructs encoded for P0 from Beet western yellow virus (pBIN61:P0 vector)(Baumberger *et al*., 2007), P1 from Rice yellow mottle virus (pCambia1300:P1Tz3 vector) (Siré *et al*., 2008) and P19 from Cymbidium ring spot virus (pBIN61:P19 vector)(Hamilton, 2002). They act on three different stages of the silencing pathway thus ensuring optimal suppression of the RNA degradation mechanism (Lacombe *et al*., 2018). All these constructions were under the control of the 35S promoter in pBin61 or pCambia vectors. These vectors were used to transform competent agrobacteria (strains C58C1 or GV3101). The transformed agrobacteria were then used for the transient transformation of *Nicotiana benthamiana* leaves.

### Agroinfiltration, plant material and experimental conditions

Strains harbouring empty pBIN61, pCambia A4:PSA, pCambia A4: RYMV_Mg1_ΔCPΔP1/ Cter PSA pCambia A4:CP_Mg1_ pBIN61:P0, pCambia1300:P1Tz3 and pBIN61:P19 vectors were separately grown overnight from precultures at 28 °C and 200 rpm in an orbital shaker using LB medium containing rifampicin (100 μg/mL) and kanamycin (50 μg/mL). The cultures were pelleted by centrifugation for 10 min at 4000 g, after which the pellets were resuspended in 10mM MgCl2 to a final OD600 of 0.5. Acetosyringone (4-hydroxy-3,5-dimethoxyacetophenone) was added to each suspension to a final concentration of 100 μM for virulence induction, and the suspensions were incubated at 24 °C for 4 h. Agroinfiltration cocktails were prepared by combining cultures for co-infiltration.

### RYMV based tool RNA accumulation assessment in *N. benthamiana* leaves

To assess the RNA accumulation of pCambia A4: RYMV_Mg1_ΔCPΔP1/ Cter PSA under RNA silencing suppression condition, pCambia A4: RYMV_Mg1_ΔCPΔP1/ Cter PSA culture combined with the cultures of silencing suppressors pBIN61:P0, pCambia1300:P1Tz3 and pBIN61:P19 following the ratio RYMV_Mg1_ΔCPΔP1/ Cter:P0:P1:P19 of 3:1:1:1 (v:v:v:v) were infiltrated into the leaves of wild-type *N. benthamiana* plants using syringes without needle. pBIN61 infiltrated following the same procedure into wild-type *N. benthamiana* plants leaves served as negative control. This first experiment also took into account the pCambia A4:PSA construct handling as described previously (pCambia A4:PSA:P0:P1:P19). 4 weeks old wild-type *N. benthamiana* plants were used. Four plants were used for the infiltration of each construct of interest and negative control. The plants were placed in a growth chamber and cultivated for 4 days before harvesting (12 h of light per day, 24 °C, 60% relative humidity). Three independent experiments were performed to generate three biological replicates for RYMV based tool RNA accumulation assessment.

### *trans* complementation of CP and/or P19

RYMV CP and/or P19 *trans* complementation has been carried out through 4 different treatments in terms of combination of constructs of interest into the leaves of wild-type *N. benthamiana* plants. These treatments consisted to independents agroinfiltration of 4 separate combinations of RYMV-based tool + CP+ P19 following the ratio 1:0,5:0,5. Effect of CP or P19 or both was assessed by removing one or the other or both constructs in the corresponding combination. To maintain the same bacterial environment, the equivalent of the removed rate (CP, P19 or both) was replaced by the corresponding rate of Pbin61 (0.5, 0.5 or 1) Four treatments were used as described in the **Table I**. Four plants were used for the infiltration of each treatments. The plants were placed in a growth chamber and cultivated for 4 days before harvesting (12 h of light per day, 24 °C, 60% relative humidity). Three independent experiments were performed to generate three biological replicates for RYMV based tool RNA accumulation assessment following each treatment through RT PCR semi quant analysis.

**Table I:** Ratio of constructs combination for agroinfiltrations.

| Treatments | Ratio of each construct per treatment |  |  |  |
| --- | --- | --- | --- | --- |
|  | RYMVΔCPΔP1/Cter PSA | CP Mg1 | P19 | Empty<br>pBIN61 |
| 1 | + (1) | - (0) | - (0) | + (1) |
| 2 | + (1) | + (0.5) | - (0) | + (0.5) |
| 3 | + (1) | - (0) | + (0.5) | + (0.5) |
| 4 | + (1) | + (0.5) | + (0.5) | - (0) |

### Determination of the optimum CP rate for the highest RYMV-based tool RNA accumulation

To determine an accurate CP rate for RYMV based tool RNA accumulation, a ratio 1 of RYMV-based tool was combined with a static rate 5 of P19 and a variable rate of CP ranging 1, 2.5 and a maximum of 4. To maintain the same bacterial environment the rate of CP removed to get different variations was replaced by the equivalent rate of the empty construct pBIN61 (**Table II**). Three different combinations of RYMV-based tool : P19 : CP : empty pBIN61 following the variation of CP rate were realized; 1 :5: 1 :3; 1 :5:2.5 :1.5; 1 : 5 :4 :0. RYMV based tool RNA accumulation under these three conditions was compare to the control combination of 1 : 0.5 ; 0.5 :0. All bacterial combinations were infiltrated into four week old *Nicothiana benthamina* leaves. Transformed plants were place in a growth chamber. 4 days later transformed leaves were harvest for RYMV based tool RNA accumulation assessment through QRT PCR analysis

**Table II:**
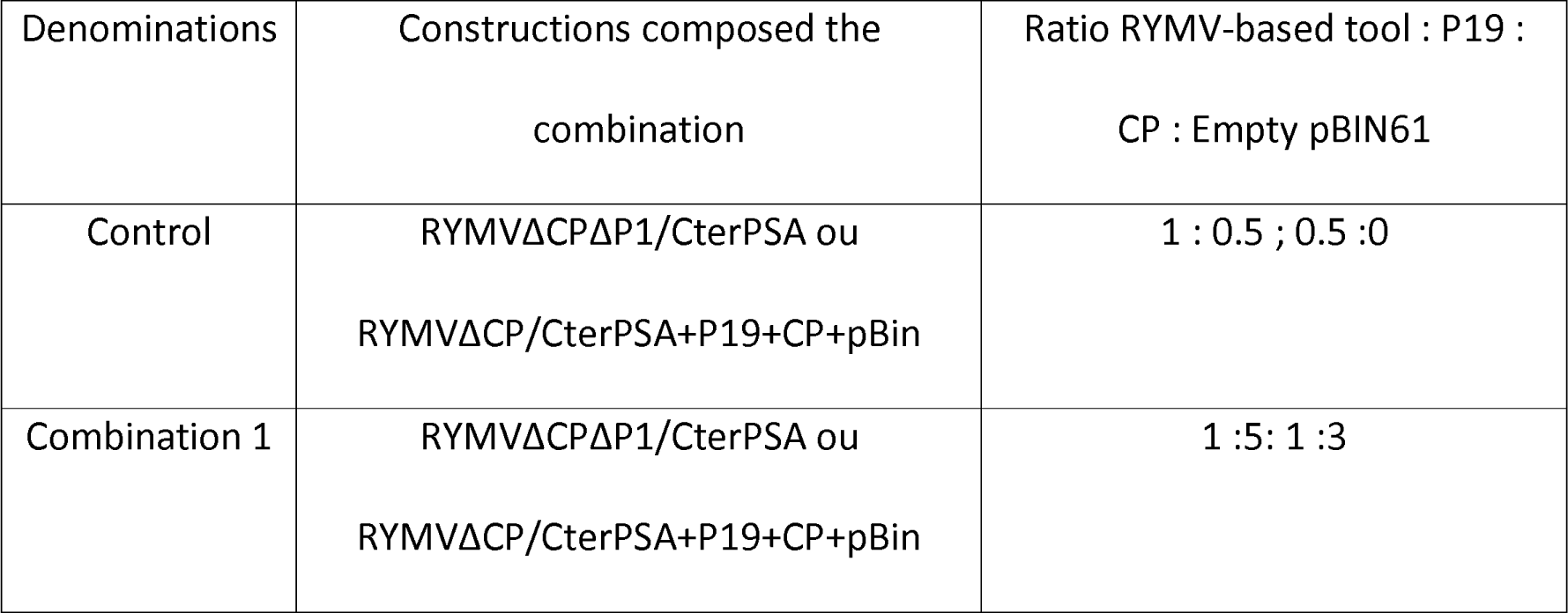
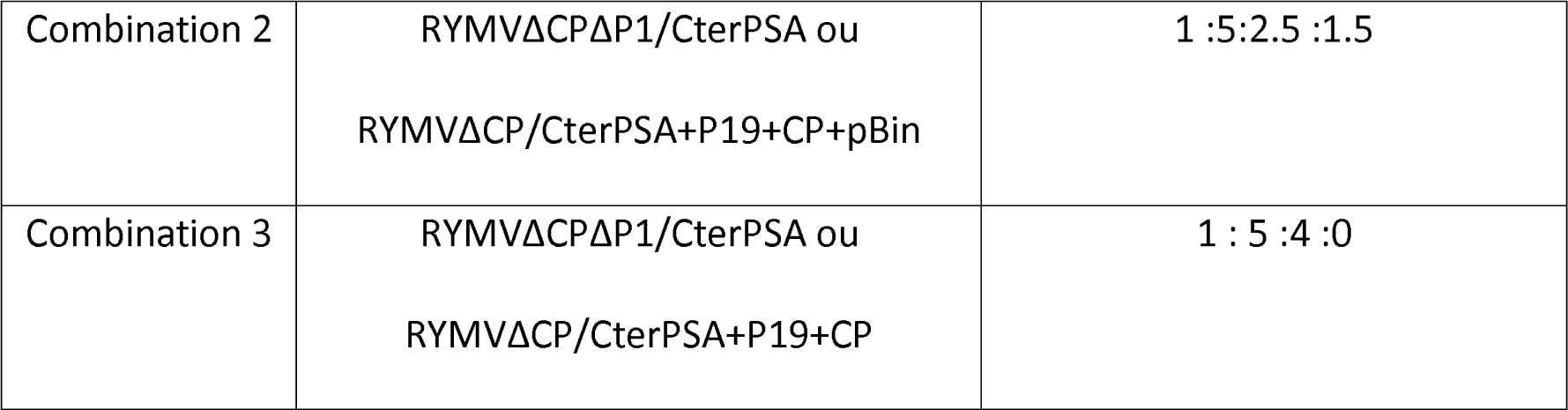
RYMV-based tool : P19 : CP : empty pBIN61.

### Semi-quantitative RT-PCR

Total RNAs were isolated from *N. benthamiana* transformed leaves using TriReagent (Sigma-Aldrich, http://www.sigmaaldrich.com). RNA samples prepared were treated with DNase according to the manufacturer’s instructions (RQ1 RNase-Free DNase from Promega), quantified with a Nanodrop spectrophotometer (Thermo Fisher Scientific, Rockford, IL, USA) and reverse transcribed into complementary DNA with Gotaq Reverse Transcriptase (Promega) using a reverse primer matching on the 3’and of viral mRNA, viral genomic and subgenomic RNA (RYMV-R 5’-CTCCCCCACCCATCCCGAGA-3’). To quantify RYMV based tool RNA accumulation, RT-PCRs were performed using primer pair RYMV-PB (5’CCAGGAAGGGCAAGAAAATC3’) and RPS-R (5’-CAGGCACGTCTTCTCGTCCGTC-3’) leading to an amplification product of 554 bp. The housekeeping gene encoding NbEIF1-α (Eukaryotic Translation Initiation Factor 1-α from *N. benthamiana*) was used as an internal control with the primer pair FNbEF1-α (5’-TCACATCAACATTGTGGTCATTGGC-3’) and RNbEF1-α (5’-TTGATCTGGTCAAGAGCCTCAAG-3’). NbEIF1-α amplification also served as loading control.

### RYMV based tool minus strain detection

For an evidence of viral vector minus strain, we performed two kind of reverse transcription on RNA from tobacco leaves transformed with pCambia A4: RYMV_Mg1_ΔCPΔP1/ Cter PSA and from those transformed with the negative control construct 35S::PSA. We used oligo dT to convert in cDNA all mRNA present in transformed cells and the forward PSA8 primer to convert in cDNA solely mRNA corresponding to viral vector minus strain, PSA8 5’-TGCTGGTGCGGACTGC-3’ is the forward primer of the couple PS8/P3 (P3 sequence: 5’CAGGCACGTCTTCTCGTCCGTC3’) amplifying a common region between the two constructs. On cDNA from the reverse transcription with Oligo dT, we conducted 2 type of PCR, one targeting the housekeeping gene encoding NbEIF1-α and another one targeting a region of RYMV ORF 2a (RYMV-2a-F 5’-GGTCGCTTTCTCACTCGCACC-3’ and RYMV-2a-R 5’-CGCAACCTTTGTGGTAGAGCG-3’) only present in pCambia A4: RYMV_Mg1_ΔCPΔP1/ Cter PSA construction. NbEIF1-α PCR served as internal control of reverse transcription and RYMV ORF 2a PCR allowed the screening of samples as an amplification should be observed on sample transformed with the viral vector and no signal for the control construct.

On cDNA from reverse transcription with PSA8, we conducted 2 types of PCRs. The first one targeted the NbEIF1-α, no signal should be detected as no plant mRNA were reverse transcribed. The second one targeted the minus strain of the viral vector, a signal corresponding to the viral vector minus strain should be detected only in sample identified as those from pCambia A4: RYMV_Mg1_ΔCPΔP1/ Cter PSA transformation. The compilation of all these PCR analysis allow to evidence whether a minus strain was present or not in samples.

### Detection of genomic and subgenomic RNA of RYMV based-tool

RNA samples prepared from *N. benthamiana* leaves as described above underwent two different reverse transcription. A reverse transcription with Oligo dT and a second reverse transcription with a mixture of Oligo dT plus a reverse primer matching on the 5’UTR region of RYMV, (RYMV-R 5’-CTCCCCCACCCATCCCGAGA-3’). Oligo dT reverse transcription target poly-A mRNA from plant cell machinery compound of plant mRNA and those of the viral vector (non genomic and non subgenomic) whereas the second reverse transcription thanks to the viral specific primer targets in addition of poly A mRNA genomic and subgenomic RNA from an eventual replication of the viral vector. A difference in PCR amplification of the viral regions between the two types of reverse transcription in favor of that performed on the cDNA from the RT Oligo dT + RYMV-R will be synonymous of an effective replication of the amplicon. Indeed, this difference will be interpreted as resulting from the detection of additional cDNA corresponding to the genomic / subgenomic non-poly A RNAs originating from the replication of the amplicon. QPCR analyzes were performed on these different cDNA with primers specific for the PSA gene of interest; PSA-F 5’CAGGCACGTCTTCTCGTCCGTC3 ’and PSA-R: 5’TGCTGGTGCGGACTGC3’. The GAPDH gene served as a normalizer gene during real-time PCR (Forward primer: GAPDH-F 5 ’GGTGTCAAGCAAGCCTCTCAC 3’, Reverse primer: 5’GATGCCAAGGGTGGAGTCAT 3 ’). The efficiency of PCR for each technical replicate was validated by LinReg software (Ruijter *et al*., 2009). The relative expression of the different genes targeted by qPCR were calculated according to the 2-ΔΔCT method (Rao *et al*., 2013). The same operation were performed on RNA from tobacco leaves transformed with the control construct 35S ::PSA. The experiment was performed three times. Comparison were done between the amounts of RNA from these two type of reverse transcription for each construct. The variation of RNA amounts between the two reverse transcription in case of the construct of interest pCambia A4: RYMV_Mg1_ΔCPΔP1/ Cter PSA was evaluated though a student t test by comparison to that of the control construct 35S ::PSA.

### pCambia A4: RYMV_Mg1_ΔCPΔP1/ Cter PSA stability analysis

We used 2 primers RYMV-PB 5’CCAGGAAGGGCAAGAAAATC3’ and RYMV-R ’CTCCCCCACCCATCCCGAGA3’ matching on the backbone of RYMV and bordering the insert for the insert stability analysis. This couple of primer allow the amplification of the fusion between insert and RYMV sequences located in region delimitated by primers. These two primer lead to an amplicon of 655 pb when the insert remains in the sequence of the viral vector. The amplicon will be smaller than 655pb if the insert is ejected. The stability of the construction was evaluated through 3 independents experiments for three different time points: j+3, j+6 and j+9.

### pCambia A4: RYMV_Mg1_ΔCPΔP1/ Cter PSA movement analysis

In tobacco plant where an important RMYV-based tool RNA were detected, three zones were defined. To check des movement of the viral vector, the infiltrated leaf was divided into 2 parts (Zones). Zone 1 represented the infiltrated area (plant cell incorporating the construct), zone 2 represented the adjacent cells close to infiltrated area, this zone was delimitated by defining 1 cm surface next to infiltrated area. The third zone was the systemic leaves. The 35S::PSA construct was used as negative control for movement evaluation, as this construct is free from any sequence of the virus. A couple of primers PSA-F 5’CAGGCACGTCTTCTCGTCCGTC3’ and PSA-R 5’TGCTGGTGCGGACTGC3’ matching on the gene of interest sequence present both in PSA entire construct and RYMV based tool, were used to probe the presence of viral vectors in these three defined areas. Three independent experiments were performed.

### Western blot

Total soluble proteins were extracted from transformed *N. benthamiana* leaves. The leaves were grounded with a mortar in liquid nitrogen until a fine powder was obtained. The powder of each sample was used to fill a 2ml Eppendorf through to 1.5 ml level, 300µl of extraction buffer (20 mM Trise HCl, 100 mM NaCl, 10 mM Na2EDTA 2H2O, 25 mM D-glucose, 0.1% Triton, 5 mM EGTA, 5% glycerol, 5mMDTT) was added to each tube. Total soluble proteins were extracted by centrifugation 30 000g for 15 min. Equal Volume of total soluble proteins were separated on a 15% (w/v) SDS polyacrylamide gel and transferred onto a Hybond-P membrane (RPN303F, GE Healthcare). Nonspecific binding was blocked with 3% (w/v) milk powder in TBS (20 mM Tris, 75 mM NaCl, and 2.5 mM MgCl2, pH 7.6) for 1h at room temperature. For detection of RYMV CP, a rabbit polyclonal antibody (Sigma Aldrich) was used at a dilution of 1:5,000 (v/v) in TBS + 3% milk powder (w/v). For PSA detection, we used a rabbit polyclonal antibody raised against *Leishmania amazonensis* excreted/secreted antigen. It has been demonstrated that this antibody displays a similar immunogenic response against *L. infantum* PSA in western experiments. This antibody was used at a dilution of 1:250 (v/v) in TBS + 3% milk powder (w/v). After washing with TBS, a peroxidase-coupled secondary antibody, anti-rabbit IgG (Immunopure Pierce Chemical, Dallas, TX, USA) at a dilution of 1:10,000 in TBS + 3% milk powder (w/v) was incubated with the membrane for 1h at room temperature. After washing with TBS, the ECL Plus Western Blotting Detection System (GE Healthcare) was used to detect both proteins. PSA expressed thanks to 35::PSA was used as a positive control for PSA detection. It was obtained from a L. infantum cDNA library. Its size is of 49 kDa. Cter PSA produced thought the pCambia A4: RYMV_Mg1_ΔCPΔP1/ Cter PSA is expressed in fusion with the remained part of RYMV CP, the fusion size is about 28 kDa.

